# Noise-induced synaptopathy impacts the long and short sensory hair cells differently in the mammalian cochlea

**DOI:** 10.1101/2023.02.27.530354

**Authors:** Yan Lu, Jing Liu, Bei Li, Haoyu Wang, Shengxiong Wang, Fangfang Wang, Hao Wu, Hua Han, Yunfeng Hua

## Abstract

In the mammalian cochlea, moderate acoustic overexposure leads to irreversible loss of ribbon-type synapse between sensory inner hair cell (IHC) and its postsynaptic auditory nerve fiber (ANF), causing a reduced dynamic range of hearing but not a permanently-elevated threshold. A prevailing view is that such ribbon loss (known as synaptopathy) selectively impacts those low-spontaneous-rate and high-threshold ANFs contacting predominantly the modiolar face of IHCs. However, the spatial pattern of synaptopathy remains scarce in the most sensitive mid-cochlear region, where long and short IHCs with distinct ribbon size gradients coexist. Here, we used volume electron microscopy to investigate noise exposure-related changes in the mouse IHCs with and without ribbon loss. Our quantifications reveal that the worst-hit areas of synaptopathy are the modiolar face of long IHCs and the pillar face of short IHCs. Moreover, we show relative enrichment of mitochondrial content in the survived ANF terminals, providing key experimental evidence for the long-proposed role of postsynaptic mitochondria in selective ribbon synapse degeneration following noise insult.

## Introduction

The mammalian cochlea decomposes sounds into different frequency components by the basilar membrane (BM) to establish a place-frequency map for resident sensory epithelial cells – inner hair cells (IHCs) – as well as their postsynaptic spiral ganglion neurons (SGNs) (for reviews, see (Fettiplace, 2017; Huet et al., 2019; Moser et al., 2020)). In rodents for instance, each IHC is contacted by about 20 predominantly unbranched peripheral dendrites of type I SGNs, forming up to 25 active zones (AZs) with characteristic electron-dense and vesicle-tethering synaptic ribbons. These specialized connections are essential for ultrafast and temporally-precise sound encoding (for reviews, see (Moser et al., 2020; Safieddine et al., 2012)). Early studies using light (Frank et al., 2010; Liberman et al., 2011; Liberman et al., 1990; Meyer et al., 2009; Wong et al., 2014) and electron microscopy (Bullen et al., 2015; Hua et al., 2021; Liberman et al., 1990; Michanski et al., 2019; Payne et al., 2021) have revealed a characteristic spatial gradient with large ribbons on the modiolar (neural) IHC face whereas small ones on the pillar (abneural) face, and such morphological diversity of presynaptic ribbons is of functional importance. As the ribbon-bound synaptic vesicle (SV) pool increases with expanding ribbon volume (Michanski et al., 2019; Payne et al., 2021; Stamataki et al., 2006), more fusion-competent SVs are made available to the synaptic release site (Joselevitch and Zenisek, 2020; Vaithianathan et al., 2016). Moreover, ribbons of different sizes are aligned to heterogeneous properties of the presynaptic Ca^2+^-channel clusters (Neef et al., 2018; Ohn et al., 2016; Ozcete and Moser, 2021; Pangrsic et al., 2015) as well as three distinct functional subtypes of SGNs (Petitpre et al., 2018; Shrestha et al., 2018; Sun et al., 2018). A prevailing view on these diversified afferent connections is to enable collective neural encoding of the entire audible range by heterogeneous presynaptic Ca^2+^-influx-release coupling and the functionally-fractionated SGNs (Heil and Peterson, 2015; Moser et al., 2020; Pangrsic et al., 2018).

However, these fine-structured afferent connections are found most susceptive to acoustic insults and non-regenerative once lost. Even moderate overexposure can lead to a permanent ribbon loss of up to ∼50% and subsequent SGN death in the rodent models (Kujawa and Liberman, 2009; Liberman et al., 2015), providing a plausible explanation for significantly declined synaptic counts observed in elder mammals (Fernandez et al., 2015; Sergeyenko et al., 2013) including human (Wu et al., 2019). As a causal consequence of ribbon loss (known as cochlear synaptopathy), coding deficits are probably implicated in weakened auditory discrimination under background noise, which is one of the hallmarks of hidden hearing loss as well as presbycusis (Kujawa and Liberman, 2015; Shi et al., 2016).

In the last decade, extensive studies on the post-exposure dynamics of cochlear synaptopathy were conducted primarily using light microscopy (LM) in different species (Fernandez et al., 2015; Hickman et al., 2020; Liberman et al., 2015; Shi et al., 2016). As the large modiolar ribbons are found to be the first undergoing degeneration following noise exposure, it turns out that the exposed IHCs often lose their prominent spatial gradient of ribbon synapses (Furman et al., 2013; Liberman et al., 2015). Although the exact mechanism of synaptopathy is still unclear, postsynaptic glutamate excitotoxicity is assumed to be an instigating factor (see reviews (Liberman, 2017; Liberman and Kujawa, 2017; Ruel et al., 2007)). In accordance with this view, the paucity of the terminal-resident mitochondria, as observed previously in the cat (Liberman, 1980), may account for the heightened vulnerability of modiolar auditory nerve fibers (ANFs, the peripheral dendrite of SGN), owing to the limited capacity of Ca^2+^ uptake during sustained synaptic transmission. However, ultrastructural quantifications of the ANF terminals in recovered ears are still missing, because the required spatial resolution to dissect these unmyelinated nerve endings can be fulfilled only by serial-section electron microscopy (EM), which is extremely labor intensive for conventional manual approach (Liberman, 2017). In addition, it has been long known that long and short IHCs coexist in the cat cochlea (Liberman, 1980). More recent studies in mice (Liu et al., 2022; Yin et al., 2014) have confirmed these two morphological subpopulations of IHCs that are prevalent in the mid-cochlear region and distinguishable based on their characteristic basolateral pole positions in an alternating arrangement. More importantly, the abneural-side type-A IHC (long) appears to have larger and more modiolar ribbons whereas the neural-side type-B IHC (short) with exclusively large ribbons gathering predominantly on the pillar face (Liu et al., 2022). This significant feature implies that two IHC subtypes may differ in their compositions of the ribbon synapses which are tuned to different physiological properties and presumably prefer one particular subtype of postsynaptic SGNs (Petitpre et al., 2018; Shrestha et al., 2018; Sun et al., 2018). Emerging questions arise about whether noise-induced synaptopathy would impact type-A and type-B IHCs differently and how.

Recent advances in the volume EM techniques have enabled the reconstruction of IHCs at unprecedented large scale and superior resolution (Bullen et al., 2015; Hua et al., 2021; Liu et al., 2022; Michanski et al., 2019; Payne et al., 2021). To address the aforementioned questions, we employed serial block-face scanning electron microscopy (SBEM) for an ultrastructural analysis on six large cochlea datasets acquired from juvenile CBA mice with or without acoustic overexposure history. This reveals mixed spatial patterns in terms of ribbon morphology in long and short IHCs following either synaptopathic or non-synaptopathic noise exposure. Besides, we verified the determinant role of mitochondrial content in the ANF survival upon noise insult.

## Results

### SBEM imaging of noise-exposed cochlear organ of Corti

In this study, we used a two-hour-long (2h) narrowband noise (8-16 kHz) to induce temporary hearing loss in male CBA mice at five weeks of age. In line with previous observations (Fernandez et al., 2015; Kujawa and Liberman, 2009; Liberman et al., 2015), upon the so-called synaptopathic (91-dB, 2h) as well as non-synaptopathic (85-dB, 2h) sounds, auditory brainstem responses (ABRs) of the exposed animals exhibit transient hearing threshold elevations with distinct frequency characters (Fig. 1A and B). Specifically, the peak threshold elevation of the non-synaptopathic group (85-dB) was at 22.6 kHz (30.00 ± 1.05 dB, n = 10 animals) on post-exposure day one (D1), while the synaptopathic group (91-dB) featured a monotonically increased threshold elevation with the frequency starting from 8 kHz and reaching a maximum at 45.2 kHz (32.86 ± 1.38 dB, n = 7 animals). On post-exposure day seven (D7), a full recovery in the ABR thresholds was observed in both groups (Fig. 1A and B). Note that there is an age-related difference in noise susceptibility between the mature (>8 weeks) and our juvenile (5-6 weeks) CBA mice (see Discussion). From the cohort, we collected cochlea tissues from six individual animals on the experimental D7 (p42, see Table 1 for more sample information). Six SBEM volumes were acquired from the mid-basal cochlear region with an estimated frequency range of 28.3 ± 0.9 kHz (Fig. 1C and S1), among which two were from animals (M1, 2) without noise-exposure history, whereas the others were from animals experiencing single exposure to either 91-dB (M3, 4) or 85-dB noise (M5, 6).

**Figure 1.**
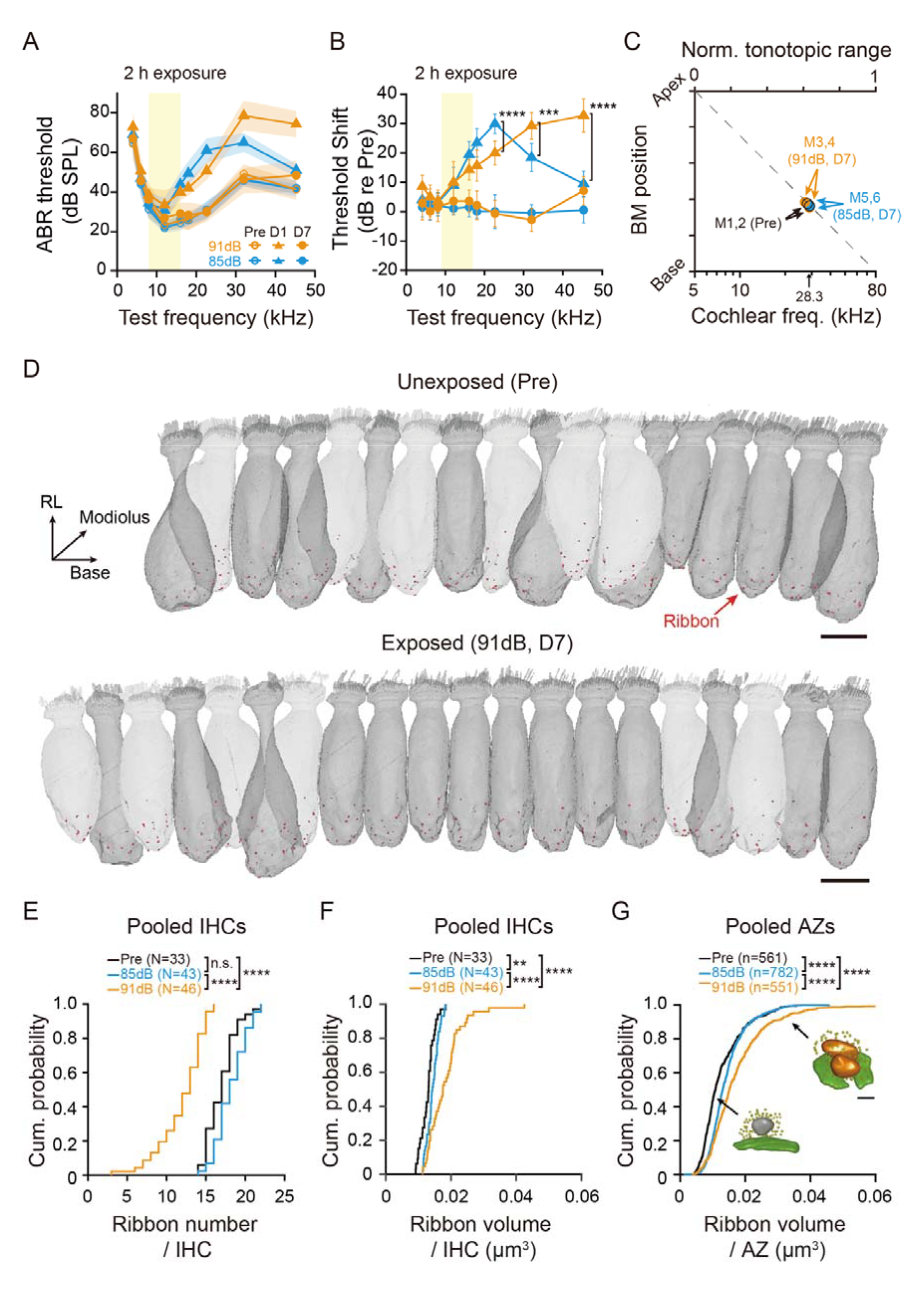
**SBEM imaging of noise-exposed cochlea.** (A) Both 85-dB (blue, n = 10 animals) and 91-dB (orange, n = 7 animals) exposures produced a transient elevation in hearing thresholds on the post-exposure day one (D1, triangles) which was fully recovered on the post-exposure day seven (D7, filled circles) in comparison with the baseline values before exposure (Pre, empty circles). (B) Relative ABR threshold shifts computed from (A). Different threshold shifts on the D1 after the 85- (blue) and the 91-dB (orange) exposures at the best-frequencies of 22.6 kHz (30.00 ± 1.05 dB vs. 20.00 ± 0.99 dB, two-sample t test, ****p < 0.0001), 32 kHz (18.50 ± 1.50 dB vs. 29.29 ± 1.09 dB, ***p = 0.0003) and 45.2 kHz (9.50 ± 1.38 dB vs. 32.86 ± 1.38 dB, ****p < 0.0001). (C) Relative BM locations determined for individual SBEM volumes (Pre: black empty circles; the 85-dB group: blue filled circles; the 91-dB group: orange filled circles), which were used to infer the best-frequencies of the SBEM-imaged IHCs based on a normalized tonotopic range from 5 to 80 kHz and yielded 28.3 ± 0.9 kHz for the mid-basal turn datasets. (D) For illustration, volume rendering of 18 reconstructed IHCs (grey) and 20 reconstructed IHCs (grey) from the unexposed (Pre) and the 91-dB exposed group (91dB, D7). Red dots at the IHC basolateral surface represent the synaptic ribbons. Dark grey and light grey reconstructed IHCs represent type-A and type-B IHCs, respectively. Scale bar, 20 μm. (E) Cumulative probability distribution of ribbon numbers from pooled IHCs of the noise-exposed (85dB and 91dB) and unexposed (Pre) ears. The median and mean ribbon numbers are 17.0 and 17.0 ± 1.9 (Pre, N = 33 IHCs), 18.0 and 18.2 ± 2.0 (85dB, N = 43 IHCs), 13.0 and 12.0 ± 2.9 (91dB, N = 46 IHCs), respectively. Two-sample Kolmogorov-Smirnov test: p = 0.1346 (Pre vs. 85dB), ****p < 0.0001 (85dB vs. 91dB), and p < 0.0001 (Pre vs. 91dB). (F) Same as (E), but with ribbon size from pooled IHCs. The median and mean ribbon volumes are 0.0131 μm^3^ and 0.0129 ± 0.0004 μm^3^ (Pre, N = 33 IHCs), 0.0143 μm^3^ and 0.0144 ± 0.0003 μm^3^ (85dB, N = 43 IHCs), 0.0175 μm^3^ and 0.0184 ± 0.0008 μm^3^ (91dB, N = 46 IHCs), respectively. Two-sample Kolmogorov-Smirnov test: p = 0.0100 (Pre vs. 85dB), p < 0.0001 (85dB vs. 91dB), and p < 0.0001 (Pre vs. 91dB). (G) Same as (E), but with ribbon size from pooled AZs. The median and mean ribbon volumes are 0.0108 μm^3^ and 0.0128 ± 0.0064 μm^3^ (Pre, n = 561 AZs), 0.0130 μm^3^ and 0.0143 ± 0.0057 μm^3^ (85dB, n = 782 AZs), 0.0151 μm^3^ and 0.0174 ± 0.0092 μm^3^ (91dB, n = 551 AZs), respectively. Scale bar, 0.2 μm. Two-sample Kolmogorov-Smirnov test: p < 0.0001 (Pre vs. 85dB), p < 0.0001 (85dB vs. 91dB), and p < 0.0001 (Pre vs. 91dB). Data in (C), (E), and (G) are presented as mean ± SD. Data in (B) and (F) are presented as mean ± SEM.

**Table 1.**
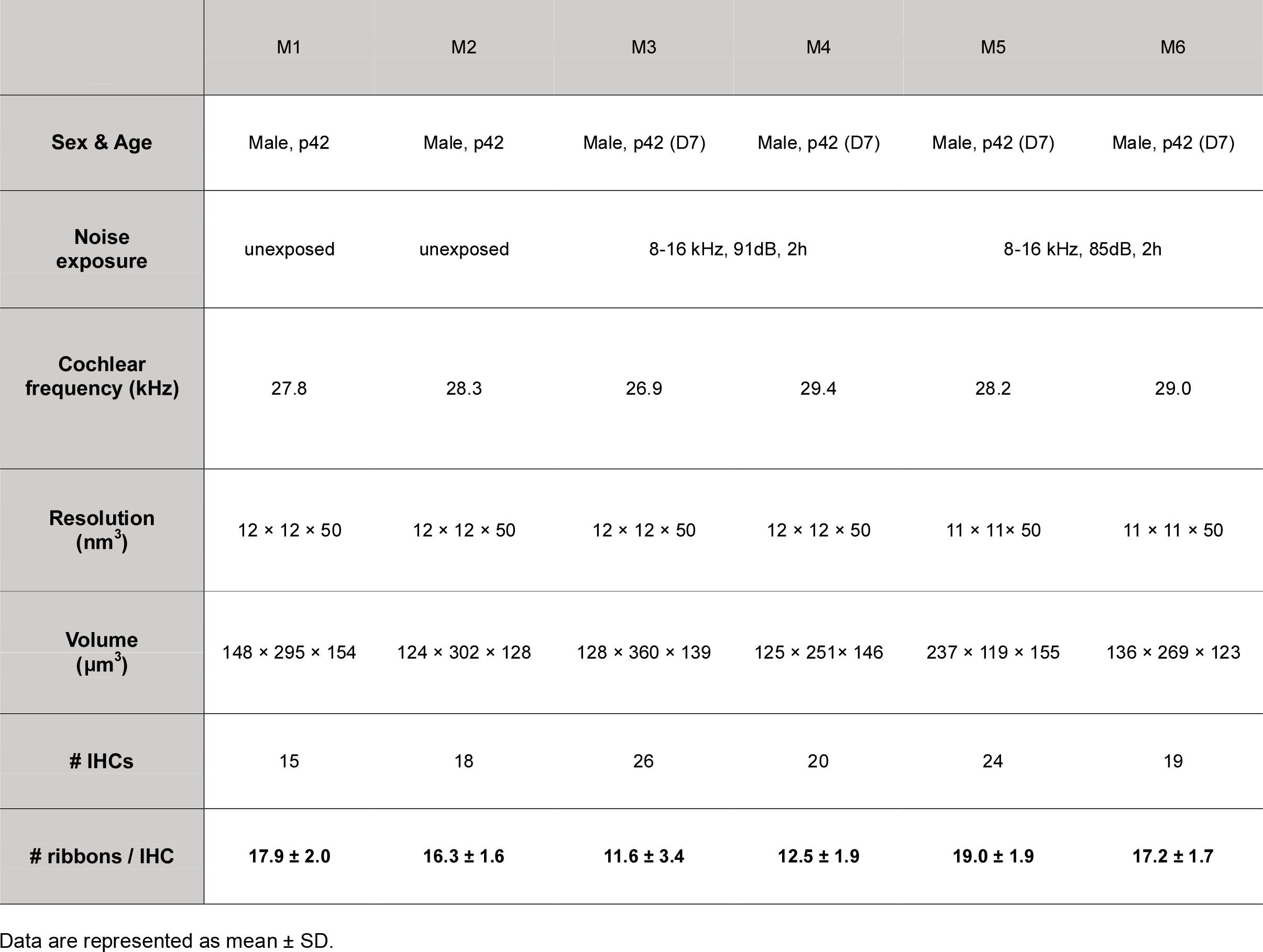
SBEM datasets from six CBA/Ca mice.

### Enlarged and multi-ribbons after acoustic overexposure

We volume reconstructed 122 intact IHCs and all 1894 ribbon synapses in the SBEM datasets (Fig. 1D and Table 1). In good agreement with prior LM studies (Fernandez et al., 2015; Grierson et al., 2022; Kujawa and Liberman, 2009; Liberman et al., 2015), a substantial ribbon loss (29.4 %) was observed in the 28 kHz cochlear region of the synaptopathic ears (Fig. 1E, 91dB: 12.0 ± 2.9, N = 46 IHCs) in comparison with the unexposed control (Pre: 17.0 ± 1.9, N = 33 IHCs, p < 0.0001) as well as the non-synaptopathic groups (85dB: 18.2 ± 2.0, N = 43 IHCs, p < 0.0001). Upon synaptopathy, the IHCs tend to have larger ribbons (Fig. 1F, 91dB: 0.0184 ± 0.0008 μm^3^, N = 46 IHCs) than those of the unexposed IHCs (Pre: 0.0129 ± 0.0004 μm^3^, N = 33 IHCs, p < 0.0001) as well as the exposed IHC without ribbon loss (85dB: 0.0144 ± 0.0003 μm^3^, N = 43 IHCs, p < 0.0001). Since we did not observe an increased occurrence of ribbon hollowing in the exposed ears (Fig. S2A), this ribbon enlargement may reflect putative activity-dependent plasticity change. Next, we compared the cumulative distribution of ribbon volume on pooled AZs and found a reduced population of single small ribbons in both exposed groups (Fig. 1G). In addition, IHCs of the synaptopathic group (91dB) appear to have a higher occurrence of multi-ribbon AZs (Fig. S2B), resulting in not only more oversized ribbons but also increased volume heterogeneity (Fig. 1G). Together, these results reveal ultrastructural insights into the adaptive plasticity of presynaptic ribbons.

### Differential synaptopathy in the long and short IHCs

All six datasets from the 28 kHz cochlear region contain both long (type-A) and short (type-B) IHCs that are staggered with alternating basolateral pole orientations (Fig. 2A). The ribbon counts suggest a comparable degree of synaptopathy across all the reconstructed type-A IHCs, meanwhile nearly half of the type-B IHCs show more profound synapse degeneration (Fig. 2B). This effect can be consistently observed from the resultant inter-IHC subtype difference in ribbon counts concomitant with the synaptopathy (Fig. S2C-H). Given that neighboring IHCs receive almost the same dose of noise exposure, this result indicates the heightened vulnerability of ribbon synapses in type-B (short) IHCs.

**Figure 2.**
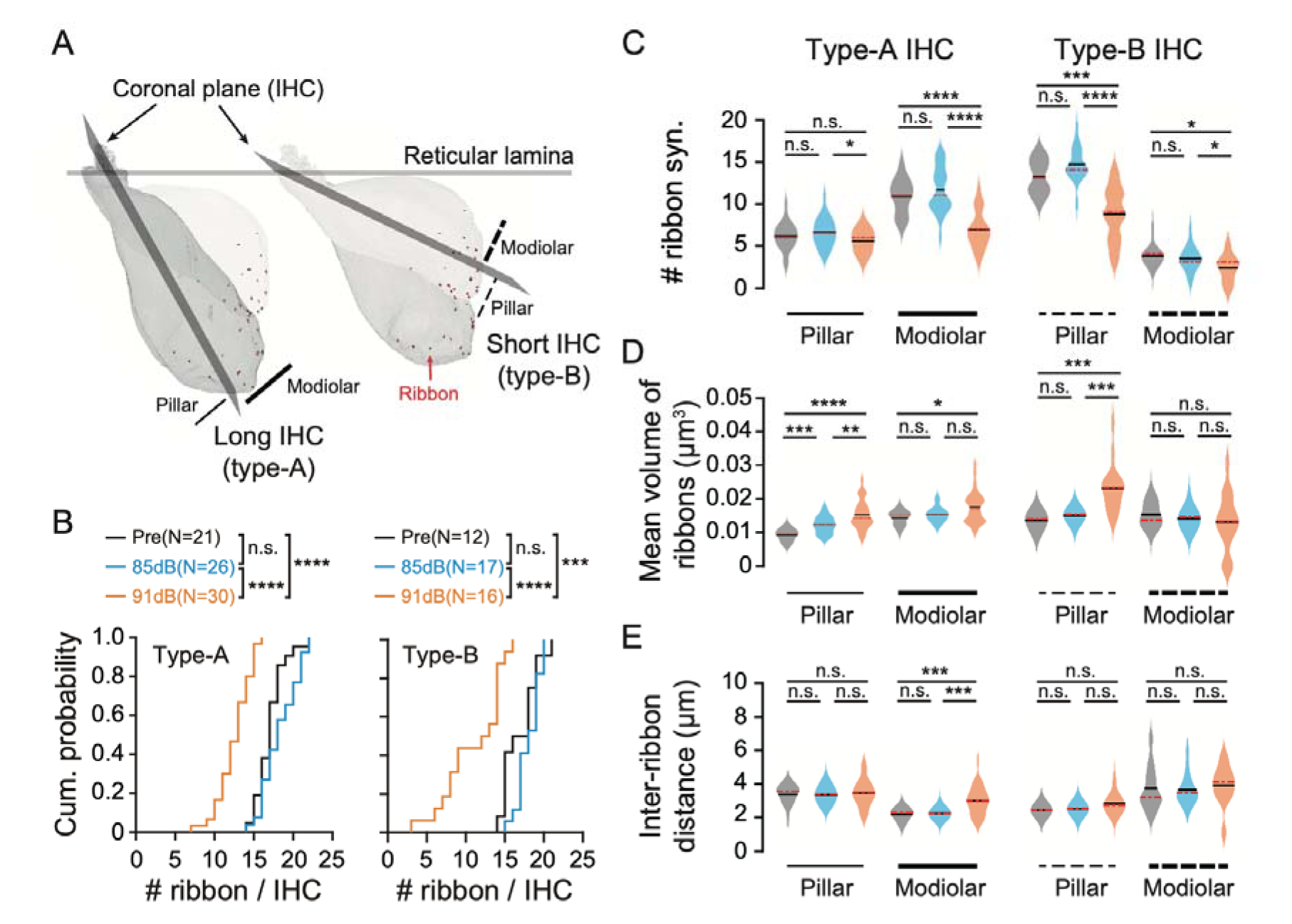
**Distinct spatial patterns of synaptopathy in the long and short IHCs.** (A) Example pairs of the type-A (dark grey) and type-B IHCs (light grey) are shown in sagittal view. They differ from each other by characteristic tilting of the coronal plane, which divides the cell body into the pillar (solid and dashed thin lines) and modiolar hemispheres (solid and dashed thick lines). Red dots at the IHC basolateral surface represent the synaptic ribbons. (B) Cumulative probability distribution of ribbon counts from pooled IHC subtypes of the noise-exposed (85dB and 91dB) and unexposed (Pre) ears. The median and mean ribbon numbers of the long IHCs (type-A) are 17.0 and 17.0 ± 1.8 (Pre, N = 21 IHCs), 18.0 and 18.3 ± 2.3 (85dB, N = 26 IHCs), 13.0 and 12.5 ± 2.1 (91dB, N = 30 IHCs), respectively. Two-sample Kolmogorov-Smirnov test: p = 0.2724 (Pre vs. 85dB), p < 0.0001 (85dB vs. 91dB), and p < 0.0001 (Pre vs. 91dB). Right: as to the short (type-B) IHCs, the median and mean ribbon numbers are 17.0 and 16.9 ± 2.2 (Pre, N = 12 IHCs), 18.0 and 18.1 ± 1.5 (85dB, N = 17 IHCs), 12.5 and 11.0 ± 3.8 (91dB, N = 16 IHCs), respectively. Two-sample Kolmogorov-Smirnov test: p = 0.1974 (Pre vs. 85dB), p < 0.0001 (85dB vs. 91dB), and p = 0.0001 (Pre vs. 91dB). (C) Left: comparisons of ribbon numbers among the type-A IHCs (solid lines) of the exposed and unexposed ears (Pre: grey, 85-dB: blue, 91-dB: orange). On the modiolar IHC face, the ribbon numbers are 10.9 ± 2.2 (Pre, N = 21 IHCs), 11.7 ± 2.7 (85dB, N = 26 IHCs) and 6.9 ± 2.3 (91dB, N = 30 IHCs). Wilcoxon rank-sum test: p = 0.3883 (Pre vs. 85dB), p < 0.0001 (85dB vs. 91dB), and p < 0.001 (Pre vs. 91dB). No significant difference among the pillar ribbons (Pre: 6.1 ± 1.6, N = 21 IHCs; 85dB: 6.6 ± 1.7, N = 26 IHCs) except for a minor decrease in the 91-dB group (91dB: 5.6 ± 1.4, N = 30 IHCs). Wilcoxon rank-sum test: p = 0.4014 (Pre vs. 85dB), p = 0.0285 (85dB vs. 91dB), and p = 0.2110 (Pre vs. 91dB). Right: same comparison for the type-B IHCs (dashed lines). Significantly fewer pillar ribbons are found in the noise-exposed IHCs (Pre: 13.2 ± 2.1, N = 12 IHCs; 85dB: 14.6 ± 2.0, N = 17 IHCs; 91dB: 8.7 ± 3.2, N = 16 IHCs). Wilcoxon rank-sum test: p = 0.1122 (Pre vs. 85dB), p < 0.0001 (85dB vs. 91dB), and p = 0.0008 (Pre vs. 91dB). Only a minor difference in the modiolar ribbons was observed (Pre: 3.8 ± 1.2, N = 12 IHCs; 85dB: 3.4 ± 1.5, N = 17 IHCs; 91dB: 2.3 ± 1.6, N = 16 IHCs). Wilcoxon rank-sum test: p = 0.4995 (Pre vs. 85dB), p = 0.0463 (85dB vs. 91dB), and p = 0.0130 (Pre vs. 91dB). (D) Comparisons of average ribbon volume in the IHCs. As to the type-A IHCs (solid lines), significant ribbon enlargement occurs on the pillar face (Pre: 0.0091 ± 0.0004 μm^3^, N = 21 IHCs; 85dB: 0.0121 ± 0.0005 μm^3^, N = 26 IHCs; 91dB: 0.0152 ± 0.0008 μm^3^, N = 30 IHCs, Wilcoxon rank-sum test: p = 0.0002 [Pre vs. 85dB], p = 0.0059 [85dB vs. 91dB], p < 0.0001 [Pre vs. 91dB]), whereas no or a minor difference among the modiolar ribbons was noted (Pre: 0.0142 ± 0.0005 μm^3^, N = 21 IHCs; 85dB: 0.0152 ± 0.0004 μm^3^, N = 26 IHCs; 91dB: 0.0174 ± 0.0009 μm^3^, N = 30 IHCs, p = 0.2895 [Pre vs. 85dB], p = 0.1600 [85dB vs. 91dB] and p = 0.0163 [Pre vs. 91dB]). As to the type-B IHCs (dashed lines), ribbon size changes are evident on the pillar IHC face (Pre: 0.0135 ± 0.0007 μm^3^, N = 12 IHCs; 85dB: 0.0151 ± 0.0004 μm^3^, N = 17 IHCs; 91dB: 0.0231 ± 0.0018 μm^3^, N = 16 IHCs, Wilcoxon rank-sum test, p = 0.0968 [Pre vs. 85dB], p = 0.0001 [85dB vs. 91dB], p = 0.0001 [Pre vs. 91dB]), but not on the modiolar face (Pre: 0.0153 ± 0.0011 μm^3^, N = 12 IHCs; 85dB: 0.0141 ± 0.0008 μm^3^, N = 17 IHCs; 91dB: 0.0130 ± 0.0021 μm^3^, N = 16 IHCs, Wilcoxon rank-sum test, p = 0.4127 [Pre vs. 85dB], p = 0.8430 [85dB vs. 91dB], p = 0.5310 [Pre vs. 91dB]). (E) Comparisons of inter-ribbon distances as a measure of local ribbon density. The modiolar face of type-A IHCs (thick solid line): 2.16 ± 0.08 μm (Pre, N = 21 IHCs), 2.22 ± 0.06 μm (85dB, N = 26 IHCs) and 2.98 ± 0.16 μm (91dB, N = 30 IHCs). Wilcoxon rank-sum test: p = 0.6922 (Pre vs. 85dB), p = 0.0004 (85dB vs. 91dB), and p = 0.0004 (Pre vs. 91dB). The pillar face of type-A IHCs (thin solid line): 3.36 ± 0.13 μm (Pre, N = 21 IHCs), 3.32 ± 0.11 μm (85dB, N = 26 IHCs) and 3.44 ± 0.15 μm (91dB, N = 30 IHCs). Wilcoxon rank-sum test: p = 0.5279 (Pre vs. 85dB), p = 0.6400 (85dB vs. 91dB), and p = 0.9470 (Pre vs. 91dB). The modiolar face of type-B IHCs (thick dashed line): 3.72 ± 0.38 μm (Pre, N = 12 IHCs), 3.62 ± 0.24 μm (85dB, N = 15 IHCs) and 3.89 ± 0.37 μm (91dB, N = 11 IHCs). Wilcoxon rank-sum test: p = 0.9417 (Pre vs. 85dB), p = 0.2540 (85dB vs. 91dB), and p = 0.4060 (Pre vs. 91dB). The pillar face of type-B IHCs (thin dashed line): 2.41 ± 0.08 μm (Pre, N = 12 IHCs), 2.48 ± 0.08 μm (85dB, N = 15 IHCs) and 2.80 ± 0.18 μm (91dB, N = 11 IHCs). Wilcoxon rank-sum test: p = 0.6430 (Pre vs. 85dB), p = 0.1320 (85dB vs. 91dB), and p = 0.0794 (Pre vs. 91dB). Data in (B) and (C) are presented as mean ± SD. Data in (D) and (E) are presented as mean ± SEM. Black and red lines represent the mean and the median, respectively.

Next, we compared the type-A and type-B IHCs in respect of their spatial pattern of synaptopathy (Fig. 2C-E). In the type-A IHCs, ribbon loss occurs exclusively on the modiolar face (Fig. 2C, Pre: 10.9 ± 2.2, N = 21 IHCs; 91dB: 6.9 ± 2.3, N = 30 IHCs, p < 0.0001), where the ANF terminals are more densely packed under normal condition. By contrast, the pillar ribbons of type-A IHCs appear noise damage-resistant (Fig. 2C, Pre: 6.1 ± 1.6, N = 21 IHCs; 91dB: 5.6 ± 1.4, N = 30 IHCs, p = 0.2110). In the case of type-B IHC, the modiolar ribbons are fewer in number but more robust to synaptopathy (Fig. 2C, Pre: 3.8 ± 1.2, N = 12 IHCs; 91dB: 2.3 ± 1.6, N = 16 IHCs, p = 0.0130) than those on the pillar side, among which degeneration of a significant portion is anticipated (Fig. 2C, Pre: 13.2 ± 2.1, N = 12 IHCs; 91-dB: 8.7 ± 3.2, N = 16 IHCs, p = 0.0008).

Intriguingly, the noise-related ribbon enlargement appears to impact the type-A IHCs on not only the modiolar face (Fig. 2D, Pre: 0.0142 ± 0.0005 μm^3^, N =21 IHCs; 91dB: 0.0174 ± 0.0009 μm^3^, N = 30 IHCs, p = 0.0163) but also the pillar face (Pre: 0.0091 ± 0.0004 μm^3^, N = 21 IHCs; 91-dB: 0.0152 ± 0.0008 μm^3^, N = 30 IHCs, p < 0.0001), where no ribbon loss was observed. Thus, the prominent ribbon size gradient is effectively compromised by eliminating large modiolar ribbons and making small pillar ribbons bigger. As to the type-B IHCs, the overweighed synaptopathy and ribbon enlargement are both found on the pillar IHC face (Fig. 2D, Pre: 0.0135 ± 0.0007 μm^3^, N = 12 IHCs; 91dB: 0.0231 ± 0.0018 μm^3^, N = 16 IHCs, p = 0.0001).

Finally, we quantified the synapse density by measuring the mean inter-ribbon distances. The result suggests that synaptopathy makes the ribbon synapses alienated from each other on the modiolar face of type-A IHC (Fig. 2E, Pre: 2.16 ± 0.08 μm, N = 21 IHCs; 91dB: 2.98 ± 0.16 μm, N = 30 IHCs, p = 0.0004) and diminishes the ribbon density gradient along the IHC modiolar-pillar axis (Pre: 2.16 ± 0.08 μm vs. 3.36 ± 0.13 μm, N = 21 IHCs, p < 0.0001; 91dB: 2.98 ± 0.16 μm vs. 3.44 ± 0.15 μm, N = 30 IHCs, p = 0.0436). Unexpectedly, in the type-B IHCs, this gradient is maintained despite a substantial ribbon loss (Fig. 2E, Pre: 3.72 ± 0.38 μm vs. 2.41 ± 0.08 μm, N = 12 IHCs, p = 0.0024; 91dB: 3.89 ± 0.37 μm vs. 2.80 ± 0.18 μm, N = 11 IHCs, p = 0.0086).

In summary, we report that the noise-related ribbon loss takes place preferentially on the modiolar side of type-A IHC and the pillar side of type-B IHC (Fig. 2C). These regions coincide with those of the highest ANF innervation density (Fig. 2E) and feature enlarged ribbons upon synaptopathy (Fig. 2D). Note that the pillar ribbons of type-A IHC seem to undergo activity-dependent enlargement as well (Fig. 2D), arguing for an IHC-wide plasticity change of ribbons in addition to local synaptic reorganization. These observations enrich our current view of noise-induced cochlear damage by revealing a set of differences in the spatial patterns of synaptopathy between two IHC morphological subtypes.

### Characterizing mitochondrial content in the ANF terminals

Prior EM study in the cat has shown that low spontaneous-rate (SR) ANFs contact exclusively the modiolar IHC face via small and mitochondrion-poor terminals (Liberman, 1980). Similarly, in our SBEM datasets of the mouse cochlea both mitochondrion-rich/poor ANF terminals can be identified (Fig. 3A). Next, we ask to what extent the postsynaptic mitochondrial content explains the mixed spatial pattern of synaptopathy in the type-A/B IHCs. For that, the numbers of mitochondria were counted manually in the ANF terminals of both unexposed and recovered ears (Fig. 3B-D). In line with early observation on the cat, type-A IHC of the rodent is innervated preferentially by mitochondrion-poor ANF terminals on the modiolar face (Fig. 3B, Pre: 7.2 ± 4.6 [pillar, n = 56] vs. 4.2 ± 3.5 [modiolar, n = 116], p < 0.0001). By contrast, the nerve terminals on type-B IHC are mitochondrion-rich and do not show a modiolar-pillar gradient (Fig. 3B, Pre: 5.9 ± 3.9 [pillar, n = 132] vs. 5.9 ± 4.1 [modiolar, n = 34], p = 0.8598). A nearly identical pattern with respect to the postsynaptic mitochondrial content was observed in the 85-dB exposed group (Fig. 3C, 85dB: 6.3 ± 4.2 [pillar, n = 144] vs. 7.1 ± 4.2 [modiolar, n = 31], p = 0.3483), suggesting limited mitochondrial plasticity upon non-synaptopathic exposure. In the case of 91-dB exposed group, we observed a diminished gradient in the mitochondrial abundance on type-A IHC (Fig. 3D, 91dB: 6.8 ± 3.7 [pillar, n = 55] vs. 5.4 ± 3.3 [modiolar, n = 80], p = 0.0321) and fewer mitochondrial-poor terminals onto type-B IHC (Fig. 3D, 91dB, 6.9 ± 3.4 [pillar, n = 91] vs. 10.0 ± 4.7 [modiolar, n = 23], p = 0.0019).

**Figure 3.**
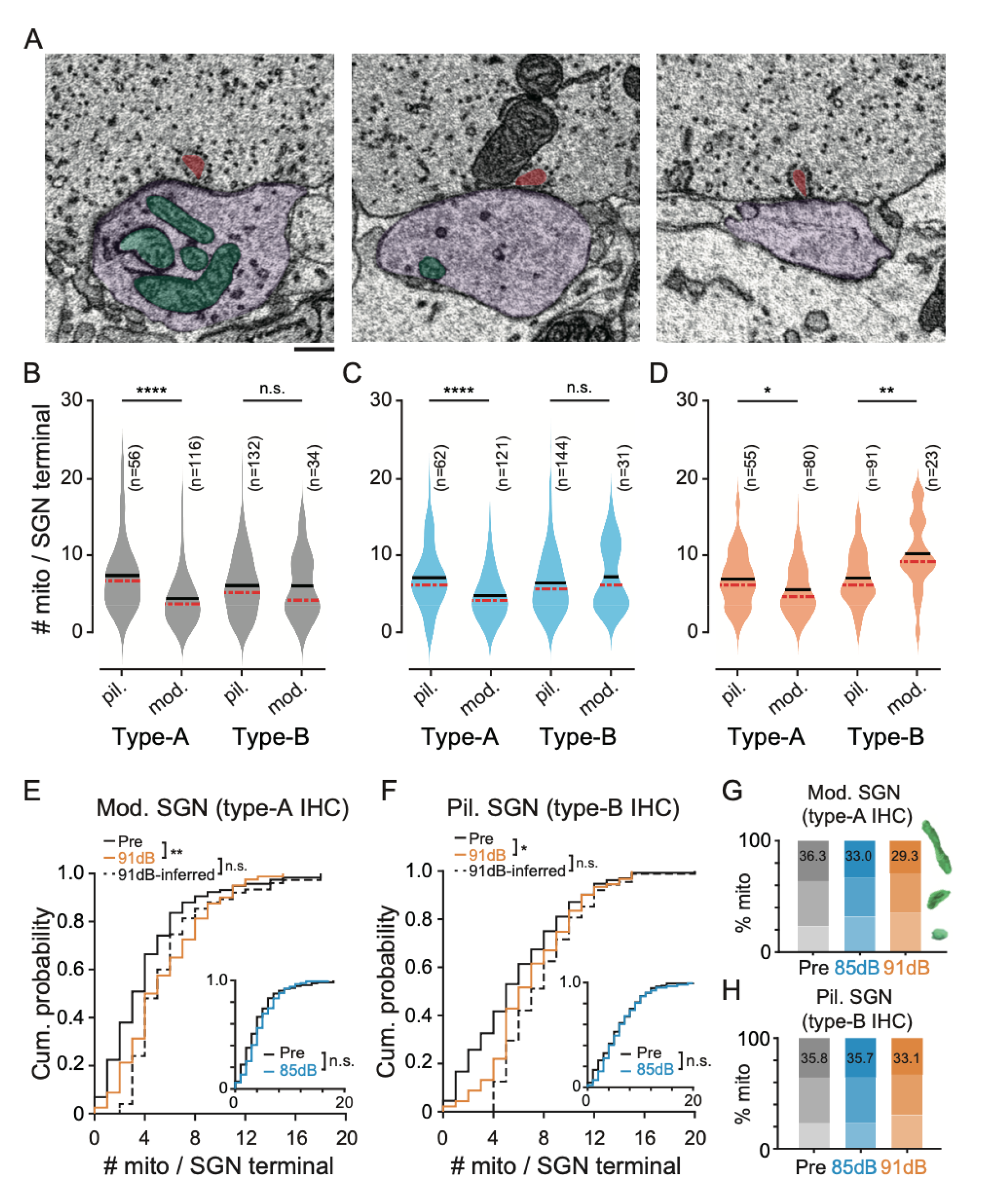
**Postsynaptic mitochondrial content in the ANF terminal.** (A) Representative EM images of different mitochondrial content (green) in the ANF terminal (purple) postsynaptic to a confirmed ribbon (red). Scale bar, 1 μm. (B) Spatial gradient of terminal mitochondrion number in the unexposed ears (grey). In the type-A IHCs, mitochondrion-rich ANF terminals are preferentially enriched on the pillar face (pil.: 7.2 ± 4.6, n = 56; mod.: 4.2 ± 3.5, n = 116, Wilcoxon rank-sum test, p < 0.0001). As to the type-B IHCs, mitochondrion-rich ANFs are prevalent on both IHC faces (pil.: 5.9 ± 3.9, n = 132; mod.: 5.9 ± 4.1, n = 34, Wilcoxon rank-sum test, p = 0.8598). (C) Same as (B), but for the 85-dB exposed group (blue). For the type-A IHCs: 7.0 ± 4.1 (pil., n = 62), 4.6 ± 3.1 (mod., n = 121), Wilcoxon rank-sum test, p < 0.0001. For the type-B IHCs: 6.3 ± 4.2 (pil., n = 144), 7.1 ± 4.2 (mod., n = 31), Wilcoxon rank-sum test, p = 0.3483. (D) Same as (B), but for the 91-dB exposed group (orange). For the type-A IHCs: 6.8 ± 3.7, (pil., n = 55), 5.4 ± 3.3, (mod., n = 80), Wilcoxon rank-sum test, p = 0.0321. For the type-B IHCs: 6.9 ± 3.4 (pil., n = 91), 10.0 ± 4.7 (mod., n = 23), Wilcoxon rank-sum test, p = 0.0019. (E) Cumulative probability distribution of mitochondrion number in the ANF terminals contacting the modiolar face of type-A IHCs. The median and mean mitochondrial numbers are 3.0 and 4.2 ± 3.6 (n = 116) for the unexposed group (black line), 4.5 and 5.4 ± 3.3 (n = 80) for the 91-dB exposed group (orange line), as well as 5.0 and 5.8 ± 3.5 (n =75) for the artificial synaptopathic group which is inferred by excluding 36% ANFs with the fewest mitochondria (black line). Inset: same distribution between the unexposed (black line) and the 85-dB exposed ears (blue line, 4.0 and 4.7 ± 3.2, n = 121). Two-sample Kolmogorov-Smirnov test: p = 0.0453 (Pre vs. 91dB), p = 0.1801 (91dB vs. 91dB-inferred), p = 0.3106 (Pre vs. 85dB). Wilcoxon rank-sum test: p = 0.0017 (Pre vs. 91dB), p = 0.5894 (91dB vs. 91dB-inferred), and p = 0.0581 (Pre vs. 85dB). (F) Same as in (E), but for the pillar ANFs of type-B IHCs. The median and mean mitochondrial numbers are 5.0 and 5.8 ± 4.0 (Pre, n = 132), 6.0 and 6.9 ± 3.5 (91dB, n = 91), 7.0 and 7.9 ± 3.2 (91dB-inferred, excluded the top 34% mitochondria-poor ANFs, n = 88), 6.0 and 6.3 ± 4.2 (inset: 85dB, n = 144), respectively. Two-sample Kolmogorov-Smirnov test: p = 0.0263 (Pre vs. 91dB), p = 0.3811 (91dB vs. 91dB-inferred), and p = 0.4157 (Pre vs. 85dB). Wilcoxon rank-sum test, p = 0.0141 (Pre vs. 91dB), p = 0.0763 (91dB vs. 91dB-inferred), and p = 0.3477 (Pre vs. 85dB). (G) The percentage of terminal mitochondria of different sizes. Comparison of ANFs contacting the modiolar face of type-A IHCs among the unexposed (Pre) and exposed groups (85dB and 91dB). Three classes of terminal mitochondria are shown (green): round (below), oval (middle), as well as large and complex (top). 36.3% (Pre), 33.0% (85dB), and 29.3% (91dB) of the total population are composed of large and complex mitochondria. (H) Same as (G), but for the pillar ANFs on the type-B IHCs. The percentages of large and complex mitochondria are 35.8% (Pre), 35.7% (85dB), and 33.1% (91dB) in the ANF terminals, respectively. Black and red lines represent the mean and the median in (B), (C), and (D).

Next, we focused on the worst-hit areas in terms of synaptopathy and compared the distribution of postsynaptic mitochondrial abundance in the unexposed and recovered ears. In the 91-dB exposed group, survived ANF terminals show significantly enriched mitochondrial content compared to those of the unexposed ear (Fig. 3E and F, 91dB: 5.4 ± 3.3, n = 80, p = 0.0017; 6.9 ± 3.5, n = 91, p = 0.0141). Such change can be somehow reproduced by a synaptopathy-mimicking exclusion of the top 35% mitochondrion-poor ANFs from the unexposed group (Fig. 3E and F, 91dB-inferred: 5.8 ± 3.5 [type-A IHC], n = 75, p = 0.5894; 7.9 ± 3.2 [type-B IHC], n = 88, p = 0.0763). Unlike the case of synaptopathy, the 85-dB exposed group does not differ from the unexposed group in terms of the ANF mitochondrial counts on either the modiolar face of type-A IHC (Fig, 3E inset, Pre: 4.2 ± 3.6, n = 116; 85dB: 4.7 ± 3.2, n = 121, p = 0.0581) or the pillar face of type-B IHC (Fig. 3F inset, Pre: 5.8 ± 4.0, n = 132; 85dB: 6.3 ± 4.2, n = 144, p = 0.3477). Further, we noted there is a reduction in the portion of large and complex mitochondria upon synaptopathy (Fig. 3G and H), suggesting upregulated mitochondrial fission in the survived ANF terminals.

## Discussion

Mechanosensory hair cells are multi-functionally compartmentalized with specialized organelles, making them attractive model systems for studying the interplay between/among subcellular components including ribbon synapses, synaptic vesicles, mitochondria and membrane cisterns (Chakrabarti et al., 2018; Esterberg et al., 2014; Hashimoto et al., 1990; Kantardzhieva et al., 2013; Lioudyno et al., 2004; Lysakowski and Goldberg, 1997; Spicer et al., 2007; Spicer et al., 1999). With the aid of advanced techniques in serial sectioning, EM reconstruction at whole-cell level has become less labor intensive, contributing to the discoveries of key morphological hallmarks in developing (Michanski et al., 2019; Payne et al., 2021), mature (Bullen et al., 2015; Dow et al., 2015; Hua et al., 2021), aging (Lauer et al., 2012), as well as noise-injured (Bullen et al., 2019) sensory organs.

SBEM is proven capable of visualizing content-rich cytoarchitecture over tens of inner and out hair cells (Hua et al., 2021; Wang et al., 2021b). Large EM volume size allows for determining the global orientation difference of the IHCs, identifying outlines of individual cell bodies in densely packed regions, as well as quantifying pre-and postsynaptic structures of the ribbon synapses simultaneously. In addition, it is known that cochlear ultrastructure can vary substantially in a tonotopy-dependent fashion, but determining the precise cochlear location of acquired EM volume has not been established yet, hampering direct inter-study comparison (Hua et al., 2021; Michanski et al., 2019; Payne et al., 2021). Recent work has demonstrated X-ray tomography as a powerful tool for the morphological study of the whole cochlea and its potential to combine with other optical methods (Keppeler et al., 2021; Lu et al., 2021). In the present work, we demonstrate the practice of targeting a given cochlear tonotopic range by using X-ray-tomographic reconstruction to guide sample trimming and sequential SBEM imaging (Fig. 1C and S1).

### Synaptopathic and non-synaptopathic acoustic exposure

Due to the non-linearity of cochlear mechanics, noise damage as well as consequent ribbon loss does not directly relate to the magnitude of ABR threshold elevation (see review (Liberman, 2017)). Thus, we calibrated the boundary condition between synaptopathic and non-synaptopathic exposures on the juvenile CBA mice (Fig. 1A and B). Although the frequency character of noise-induced ABR threshold shifts is consistent with those previously seen on the mature animals, our juvenile mice have acquired ribbon loss already upon a lower noise dose (91 dB, 2h), which has been reported as a non-synaptopathy-inducing condition for the animal at 16 weeks of age (Fernandez et al., 2015). Moreover, complementary to an early study (Kujawa and Liberman, 2006) showing accelerated hair cell loss in the young-exposed (4-8 weeks) but not in the old-exposed (≥ 16 weeks) animals to a 100-dB noise exposure, this result corroborates that the juveniles are more vulnerable to acoustic overexposures in terms of not only neuropathy but also synaptopathy. As permissible noise levels for occupational exposure are established in adults (Daniell et al., 2006; Davis and Clavier, 2017), it may be advisable to take preventive actions with stricter guidelines in schools.

### Ribbon loss and plasticity in the mature IHCs

It has been shown that ribbon size is of functional significance by determining the pool of synaptic vesicles available for fusion and tightly regulated by presynaptic calcium channels (Frank et al., 2009; Ozcete and Moser, 2021; Sheets et al., 2012). Emerging volume EM techniques have been employed to dissect the morphological maturation of synaptic ribbons, ranging from late embryonic to adult stages (Hua et al., 2021; Michanski et al., 2019; Payne et al., 2021). In the mature IHCs however, evidence for ribbon maintenance and activity-dependent plasticity remains scarce (Voorn and Vogl, 2020). Consistent with prior observations using LM (Liberman et al., 2015), post-exposure enlargement of synaptic ribbons was verified by quantitative SBEM (Fig. 1F and G). More importantly, the enlarged ribbons are not associated with “hollowing” cores or fewer tethered vesicles, arguing against a perturbed ribbon function beyond one week after the noise exposure (Fig. S2A). Besides, it is unexpected to find multi-ribbon AZs more prevalent in the synaptopathic ears (Fig. 1G and S2B), because enhanced synaptic transmission may lead to heightened glutamate-mediated excitotoxicity to the ANF terminals as one would commonly anticipate. Therefore, the functional property of this structural variant remains to be explored, given that the multi-ribbon AZ is not only retained into adulthood (Hua et al., 2021; Michanski et al., 2019) but also a hallmark of cochlear aging (Stamataki et al., 2006). It has been recently proposed that such molecular bypass of strict constraint on the maximum ribbon size may allow extra synaptic strengthening to maintain overall neurotransmission level with reduced afferent contacts (Michanski et al., 2019). Finally, the appearance of enlarged and multi-ribbons leads to an increased ribbon size heterogeneity (Fig. 1G), implying that the IHC coding deficit might not be a simple consequence of selectively eliminating large ribbons and possibly involve maladaptive changes in both pre-and postsynaptic structures.

### Distinct spatial pattern of synaptopathy in the long and short IHCs

In the middle cochlear region, staggered IHCs with alternating basolateral pole positions are prevalent (Yin et al., 2014). It has been recently shown that type-A IHC (abneural side, long) has larger and more ribbons on the modiolar face, whereas type-B IHC (neural side, short) contains exclusively large ribbons but predominantly on the pillar face (Liu et al., 2022). As the postsynaptic SGNs with molecular diversity are arranged with distinct spatial preferences (Petitpre et al., 2018; Sherrill et al., 2019; Shrestha et al., 2018; Sun et al., 2018), it has been proposed that distinct ribbon gradients with a limited overlap of these two morphological subtypes may introduce disproportionated SGN compositions, thereby creating different IHC receptive fields to encode over a broad audible range (Liu et al., 2022).

It has been found in various animal models that the low-SR ANFs face predominantly large modiolar ribbons (Liberman, 1982) and are more vulnerable to noise insult (Furman et al., 2013). This notion is further strengthened by the observation of preferential ribbon loss on the modular IHC face in the recovered ears (Liberman et al., 2015). Our results suggest that the worst-hit areas of noise damage are the IHC subregions with high ribbon density, namely the modiolar face of type-A as well as the pillar face of type-B IHCs (Fig. 2C). Different ways of quantification for LM and SBEM may explain this discrepancy. In the LM studies (Hickman et al., 2020; Liberman et al., 2015; Yin et al., 2014), a global modiolar-pillar axis is routinely used and thereby the type-A IHC contributes more to the LM-classified “pillar” ribbons while the type-B IHC to the “modiolar” ribbons. Indeed, we find there is severer ribbon loss in the exposed type-B IHC (Fig. 2B), matching the modiolar preference. But in fact, the type-B IHC loses more ribbons on the pillar face as uncovered by our SBEM quantification (Fig. 2C).

The mixed spatial patterns of synaptopathy, accompanied by substantial ribbon volume changes, may reflex some sort of pre-and post-synaptic adaptation and reorganization. In line with this notion, a recent physiology study has revealed a post-exposure reduction in the ANFs of both low and high SR in mice (Suthakar and Liberman, 2021). It is attempting to speculate that mature ribbon synapse also undergoes activity-dependent plasticity (Voorn and Vogl, 2020), presumably under tight regulation of local interplay between presynaptic-and mitochondrial-Ca^2+^ influx, as shown recently in the zebrafish lateral-line hair cells (Lukasz et al., 2022; Wong et al., 2019). If so, there might be an underdetermined role of proper presynaptic organization, for instance, the mitochondrial and membrane networks (Bullen et al., 2015; Liu et al., 2022; McQuate et al., 2023), in shaping the spatial gradient of IHC ribbons to match preferred postsynaptic SGN targets for a normal hearing function. In turn, noise-induced presynaptic disorganization, as reported recently (Bullen et al., 2019), may lead to limited recovery and maladaptation with respect to the synaptic ribbon diversification and thereby an impaired level encoding of sounds.

### Postsynaptic mitochondrial content of the ribbon synapse

Despite no direct proof, it is a common belief that the paucity of terminal mitochondria causes heightened vulnerability of those ANFs postsynaptic to the modiolar ribbons by glutamate-mediated calcium overload (Liberman, 2017). This notion remains speculative because fewer mitochondria in the low-SR ANF terminal are evident so far only in the cat (Liberman, 1980, 1982) and noise-induced synaptopathy has been suggested not as selective as originally expected (Chen et al., 2019). Here, our result provides critical pieces of experimental evidence from the more widely used rodent model (Fig. 3). Unlike the extreme scenario in the cat cochlea, we found that mitochondrion-rich and -poor ANF terminals of mice are not clearly segregated along the modiolar-pillar axis. Nonetheless, the remaining ANF terminals of the synaptopathic ears demonstrate an upregulated mitochondrial content, matching the inference based on survivorship bias (Fig. 3E and F). As mitochondrion-poor terminals are prevalent at the intermediate zone between two staggered IHCs rather than the outer edges of the pair (Fig. 2C and 3B), we speculate that local high synapse density may lead to an overcrowded postsynaptic space, which favors the small-sized ANF terminal with limited mitochondrial content.

Note that we do capture a considerable fraction of big and mitochondrion-rich ANF terminals facing large presynaptic ribbons in those regions (Fig. 3E and F), consistent with the most recent LM quantification by CTBP2 and GluA2 staining in mice (Reijntjes et al., 2020). Nevertheless, it remains elusive for future investigations how this uneven ribbon synapse distribution along the IHC basolateral pole and heterogenous mitochondrial content in putative low-and high-SR ANF terminals are modulated by the transcriptomic changes in response to sensory experience including acoustic trauma (Frank et al., 2023; Milon et al., 2021).

## Materials and Methods

### Animals

In this study, CBA/Ca mice (male, p28) were purchased from Shanghai Jihui Experimental Animal Feeding Co., Ltd. (Shanghai, China). During the experiment (except for the noise exposure), animals were raised in the same acoustic environment and sacrificed at the same age (p42). Experiments were conducted at the Ear Institute of Shanghai Jiao Tong University School of Medicine and Shanghai Institute of Precision Medicine. All procedures were approved by the Institutional Authority for Laboratory Animal Care of Shanghai Ninth People’s Hospital (SH9H-2020-A65-1).

### Acoustic overexposure

The acoustic overexposure was carried out in a sound-attenuating chamber (Shanghai Shino Acoustic Equipment Co. Ltd, China). Two acoustic overstimulation protocols were used in this study: an octave-band of noise (8-16 kHz) at 91 or 85 dB for 2 hours (Fig. 1B). During the noise exposure, awake animals were kept without restraint in 18 x 18 x 10 cm^3^ sized cells (1 animal/cell) in a subdivided cage by barbed wires. The cage was placed about directly below the horn of the loudspeaker (X4, HiVi Acoustic, Inc.) connected to a sound card (US-366, TASCAM, Inc.). The octave-band waveform was custom-made (a gift from Prof. Bo Zhong, National Institute of Metrology, China) and real-time monitored by Adobe Audition Software. Before each exposure session, the noise was calibrated to the target sound pressure level using an acoustimeter (AWA6228+, Hangzhou Aihua Instruments Co., Ltd., China) placed in the cage.

### Auditory Brainstem Responses (ABRs)

ABR measurements were conducted as previously described (Wang et al., 2021a; Zhang et al., 2022). In brief, animals were anesthetized, and the body temperature was maintained near 37°C using a regulated heating pad with a thermal probe placed under the abdomen (Homeothermic Monitoring System, Harvard Apparatus, US). Sound stimuli, which were generated by SigGen RP (Tucker-Davis Tech. Inc., US), were delivered via a speaker (MF1, Tucker-Davis Tech. Inc., US) placed 10 cm away in front of the animal’s vertex. ABRs were recorded via three subdermal needle electrodes placed at the animal’s vertex (active electrode), left infra-auricular mastoid (reference electrode), and right shoulder region (ground electrode). Signal digitalization and acquisition were done with the software BioSigRZ (Tucker-Davis Tech. Inc., US). The raw signals were amplified (5000x) and bandpass filtered (0.03-5 kHz). The sound level of stimulus started from 90 dB SPL and was decremented in 5 dB steps to ∼10 dB below a threshold, which was defined as the lowest stimulus needed for evoking visible responses.

### Sample Preparation

*En bloc* EM staining of the cochlea was performed following the established protocol (Lu et al., 2021). In brief, after being anesthetized with 2% isoflurane inhalation, animals were decapitated, and temporal bones were dissected from both sides. Under the microscope, the cochleae were immediately perfused through the round and oval windows with an ice-cold fixative mixture containing 0.08 M cacodylate (pH 7.4, Sigma-Aldrich, US), 2% paraformaldehyde (Sigma-Aldrich, US) and 2.5% glutaraldehyde (Sigma-Aldrich, US). Post-fixation and sequential decalcification were carried out at 4°C by immersing in the same fixative and with the addition of 5% EDTA (Sigma-Aldrich, US) for 5 hours each. The decalcified cochleae were then washed twice with 0.15M cacodylate buffer (pH 7.4) for 30 min each, and then immersed in 2% OsO_4_ (Ted Pella, US), 2.5% ferrocyanide (Sigma-Aldrich, US), and 2% OsO_4_ at room temperature for 2, 2, and 1.5 hours, respectively. All staining solutions were buffered with 0.15M cacodylate buffer (pH 7.4). After being washed with 0.15M cacodylate buffer and nanopore-filtered water for 30 min each, the cochleae were sequentially incubated in 1% freshly made and filtered thiocarbonhydrazide solution (saturated aqueous solution, Sigma-Aldrich, US), 2% OsO_4_ aqueous solution and lead aspartate solution (0.03M, pH 5.0, adjusted by KOH) at 50°C for 1, 2 and 2 hours, with immediate washing steps. For embedding, the cochleae were dehydrated through a graded acetone-water mixture (50%, 75%, 90% acetone at 4°C for 30 min each) into pure acetone (3 times at room temperature for 30 min each). Then the dehydrated cochleae were infiltrated with a 1:1 mixture of acetone and Spurr resin monomer (4.1g ERL 4221, 0.95g DER 736, 5.9g NSA and 1% DMAE; Sigma-Aldrich, US) at room temperature for 6 hours on a rotator, followed by 1:2 mixture of acetone and resin monomer at room temperature for overnight on a rotator. Infiltrated cochleae were then incubated in pure resin for 8-12 hours before being placed in embedding molds (Polyscience, Germany) and incubated in a pre-warmed oven at 70°C for at least 72 hours.

### Correlative X-ray Microscopy (XRM) and Serial Block-face Electron Microscopy (SBEM)

For high-resolution XRM imaging of the cochlea, excess resin around the cochlea was carefully removed using a trimmer (TRIM2, Leica, Germany) equipped with a diamond knife head (#227172, Anton Meyer & Co. Ltd). Cochlea samples were mounted upright along the conical center axis on a metal rivet (3VMRS12, Gatan, UK) and scanned with an XRM (Xradia 520 Versa, Carl Zeiss, Germany) at a pixel size of < 4.7 µm, source voltage 60 kV, power 5 W and exposure time per radiograph 1 sec. Cochlear 3D reconstructions based on acquired XRM datasets were visualized and further analyzed in 3D image processing software (AMIRA, Thermo Scientific, US). The whole extent of the basilar membrane was annotated with marked segments of interest using a published cochlear place-frequency map of the mouse as a reference (Muller et al., 2005). After having trimmed the cochleae coronally to the targeted location, we imaged (30 × 30 nm^2^, 60 × 60 μm^2^ per image) the block face of each sample using an SEM (Gemini300, Carl Zeiss, Germany) for *post hoc* confirmation of the frequency range of the SBEM volumes.

The samples were remounted with the previously SEM-scanned block face on top and further trimmed down to a block size of about 800 × 800 x 500 μm^3^ (xyz). Serial sections were imaged using a field-emission SEM (Gemini300, Carl Zeiss) equipped with an in-chamber ultramicrotome (3ViewXP, Gatan) and back-scattered electron detector (Onpoint, Gatan). In total, six SBEM datasets were analyzed in this study (Fig. 1C), two (M1 and M2) unexposed cochleae, as well as four (M3-6) noise-exposed cochleae. Imaging parameters were as follows: incident electron beam energy, 2 keV; pixel dwell time, 1.5 - 2.0 μs; pixel size, 11 - 12 nm; cutting thickness 50 nm; focal charge compensation, 100%; vacuum chamber pressure, ∼2.8×10^-3^ mbar. Consecutively acquired slices were aligned offline using a cross-correlation function in MATLAB (MathWorks, US) and cubed for volume tracing in webKNOSSOS (Boergens et al., 2017; Hua et al., 2021).

### Reconstruction of IHC, Mitochondrion, and Ribbon Synapse

Using the previously trained 3D U-net models (Liu et al., 2022), the in-plane probability maps of IHCs were generated by running the networks with a blocking scheme. In brief, a distance-transform-watershed algorithm was applied to each thresholded probability map to produce over-segmented super-pixels for IHCs. To achieve volume reconstruction, we used a 3D connection algorithm to connect the 2D super-pixels and output the reconstructed 3D objects. From all the datasets, 122 IHCs were volume reconstructed. Manual proofreading and correction were done in webKNOSSOS (Boergens et al., 2017) on the auto-segmentation results of ribbon synapses. The coronal plane of each IHC was determined by two lines and its center point. The first line was parallel to one IHC row which was computed by fitting a line with IHC centers on it. Another line was the principal axis of each IHC computed by principal component analysis (PCA). Each IHC cell body was then divided into pillar and modiolar hemispheres based on the coronal plane and with individual ribbon synapses assigned objectively to either hemisphere.

### Statistical Analysis

All data analysis and statistical tests were carried out using self-written scripts and built-in functions in MATLAB (release 2021a) and the Statistics Toolbox (MathWorks, Inc., US). The comparisons between groups were done using the two-sample t test (*ttest2*) for Fig 1B, S2A-B. The distribution comparisons between groups were done using two-sample Kolmogorov-Smirnov tests (*kstest2*) for Figure 1E-G, 2B, 3E-F. The distribution comparisons between groups were done using the Wilcoxon rank-sum test (*rank-sum*) for Figure 2C-E, 3B-F. The significance level of statistical tests was denoted as n.s. for p > 0.05, * for p < 0.05, ** for p < 0.01, *** for p < 0.001, and **** for p < 0.0001.

## Author Contributions

Y.H. designed and supervised the study; Y.L. and B.L. carried out the animal experiments, Y.L., J.L., H.W., and S.W. analyzed the data; J.L. and H.H. contributed to the segmentation and 3D reconstruction; F.W. assisted with the EM acquisition; Y.H., Y.L., H.W., and J.L. drafted the manuscript. All authors commented on the manuscript.

## Acknowledgments

We thank X.D. for the help in the early stage of this work and thank Shanghai Jiao Tong University Instrumental Analysis Center for the assistance of XRM imaging. This work was supported by the National Natural Science Foundation of China (82171133 to Y.H., 32171461 to H.H.), Industrial Support Fund of Huangpu District in Shanghai (XK2019011 to Y.H.), Innovative Research Team of High-Level Local Universities in Shanghai (SHSMU-ZLCX20211700), the Shanghai Key Laboratory of Translational Medicine on Ear and Nose Diseases (14DZ2260300).

